# Evolution of the Telencephalon Anterior-Posterior Patterning by Core Endogenous Network Bifurcation

**DOI:** 10.1101/2023.03.30.534890

**Authors:** Chen Sun, Mengchao Yao, Ruiqi Xiong, Yang Su, Binglin Zhu, Ping Ao

## Abstract

How did the complex structure of telencephalon evolve? Existing explanations are based on phenomena and lack the first principle. The Darwinian dynamics and the endogenous network theory established a few years ago provide a mathematical and theoretical framework of a dynamical structure, and a general constitutive structure for theory-experiment coupling, respectively, for answering this question from the first principle perspective. By revisiting a gene network that explains the anterior-posterior patterning of the vertebrate telencephalon, we found that with the increase of the cooperative effect in this network, the fixed points gradually evolve, accompanied by the occurrence of two bifurcations. The dynamic behavior of this network consists with the knowledge obtained from experiments on telencephalon evolution. Furtherly, our work drew an answer quantitatively of how the telencephalon anterior-posterior patterning evolved from the pre-vertebrate chordate to the vertebrate and gave a series of verifiable predictions in a first principle manner.

**Figure Abstract:** 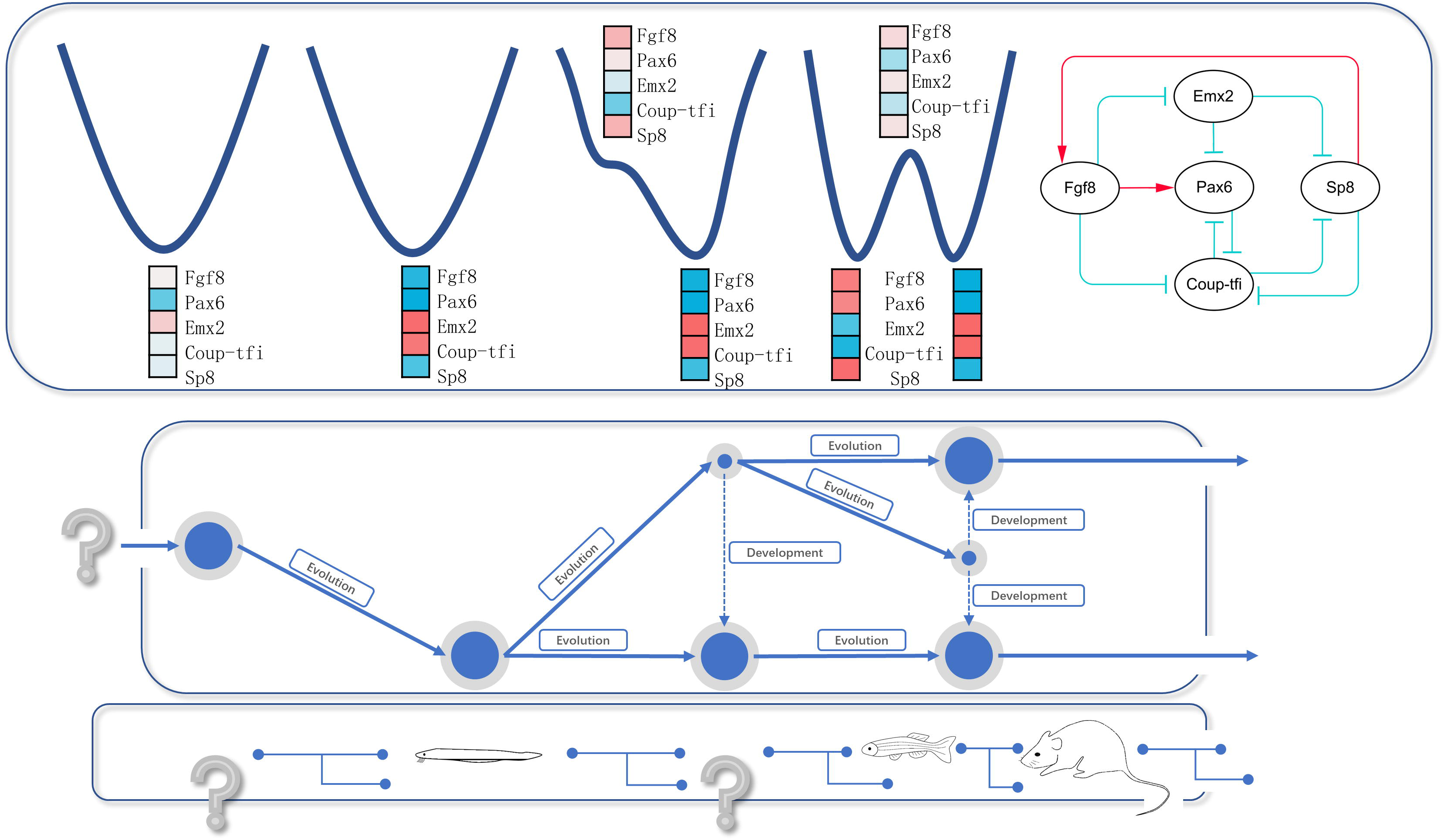

## Introduction

As the material basis of cognitive abilities, the telencephalon results from hundreds of millions of years of evolution(Holland 2015; Briscoe and Ragsdale 2019; Chakraborty and Jarvis 2015). A series of events were discovered, like genome duplication and new genes emergence(Kaessmann 2010; Putnam et al. 2008), core gene network interconnectivity increasing(Gil-Gálvez et al. 2022), and neuropathway duplication(Chakraborty and Jarvis 2015), providing phenomenological explanations for how the complex structure of the telencephalon has evolved to this day. But like research on the celestial motion before the introduction of Newtonian mechanics, these phenomena-based explanations lack the fundamental core: the first principle.

Recently, some researchers also worried that researches in life science pay more attention on the experiment itself than the theory-experiment coupling(MacArthur 2023; Smaldino 2019; Nurse 2021); or even had a circling back sensation, at least in cancer research, after half a century of the journey(Weinberg 2014). Indeed, the phenomena of life are full of complexity, and the complexity science itself is still a newborn one. The discovery of a first principle in life is bound to be difficult. Nonetheless, there were fruits born in this “circle”. Decades after the introduction of Boolean dynamics into the gene network(Kauffman 1969; Toulouse et al. 2005), the analysis of the dynamical structure of the hydrogen model in life science: the phage-λ gene switch(Zhu et al. 2004), provided an example of application of a general law alone with the mathematical framework for the evolution dynamics(Ping Ao 2005; Kwon, Ao, and Thouless 2005). Following this footpath, a theory provides the general constitutive structures of evolution dynamics on the gene network level, the endogenous network theory(Ping Ao et al. 2008) was established years after the postulation of the system biology(Hood 2003), unifies the gene network dynamics and the landscape concept raised by Wright and Waddington, allowing the qualitative and quantitative comparisons between theories and experiments(Ping Ao 2021; Prokop 2022). Under these theoretical sets, a series of efforts were made to address questions on cancer and development(Yuan et al. 2017). Now we may have enough mathematical and theoretical framework as well as the knowledge obtained from experiments to address the question of how our brain evolved at the gene network dynamics level in a first principle way.

In this work, the attempt was made by revisiting a pre-constructed model(Giacomantonio and Goodhill 2010; 2014; Goodhill 2018) of telencephalon anterior-posterior patterning in vertebrates containing 5 nodes and 10 edges, demonstrated that the dynamic behavior of such a relatively small network is necessary to draw an answer under the mathematical and theoretical framework mentioned above.

## Results

### A coarse-grained core endogenous network for telencephalon already exists

A Boolean 5-node network has been constructed, which sufficiently reproduced the experimentally observed gradients of anterior-posterior patterning of early cortical development in mammals(Giacomantonio and Goodhill 2010; Goodhill 2018). From the perspective of data fitting, the existing experimental data were not sufficient to uniquely specify interactions of a base network. And if such a base network exists, how the latent factors regulate the base network is not clear(Giacomantonio and Goodhill 2010; 2014; Goodhill 2018).

But in the other way, Giacomantonio and Goodhill have constructed a network with robust dynamic behavior, which is exactly one of the most important features of the “backbone”, or say, the “core” structure of an endogenous network shaped by evolution that we argued in 2008, one of the others is being conserved among species during evolution(Ping Ao et al. 2008; Yuan et al. 2017). We found ortholog exists in amphioxus, one of the closest living vertebrate relatives that diverged from the vertebrate lineage around 550 million years ago, of all 5 genes(Putnam et al. 2008; Marlétaz et al. 2018; Takatori et al. 2008) in this network. Nevertheless, Benito-Gutiérrez *et al*. discovered the resemblance of the expression pattern between mature amphioxus cerebral vesicle and the developing vertebrate telencephalon(Benito-Gutiérrez et al. 2021). These facts indicate that the network Giacomantonio and Goodhill constructed is conserved in the pre-vertebrate chordate and the vertebrate during evolution.

To further confirm the robustness of the model, we generated 128 Boolean function variations based on the network (Figure 1. A) described by Goodhill in 2018(Goodhill 2018) by using the combination of replacing “AND” with “OR”. With the default operators, 2 point attractors which correspond to the expression pattern of the posterior (boole_p1) or anterior (boole_p2) telencephalon separately and 2 linear attractors (boole_li3 and boole_li14) were generated by exhausting all initial vectors (Figure 1. B). For all Boolean variations, we obtained 26 attractors with 4 point ones and 22 linear ones (Figure 1. B). Among them, two attractors, the posterior-like attractor boole_p1 and the anterior-like attractor boole_p2, were obtained in all Boolean variations (Figure 1. C, E). The size of the attraction basins with each attractor in all variations, in which the boole_p1 and the boole_p2 was equivalently the most common attractor among all (Figure 1. D, E). These results further confirmed the robustness of the Boolean dynamic behavior of this network.

**Figure 1.**
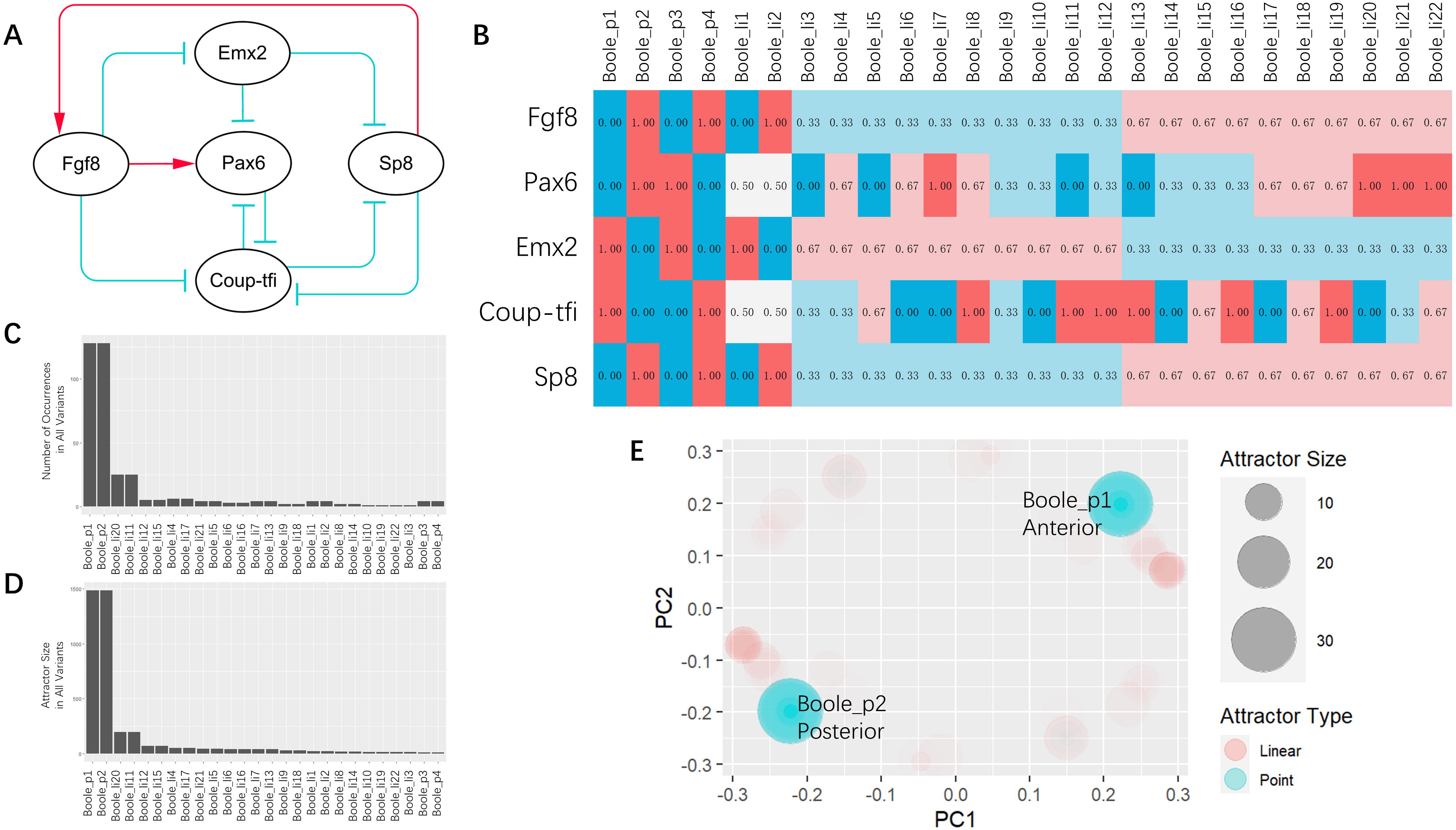
Boolean dynamics of the core endogenous network for telencephalon anterior-posterior patterning. A, The network we used in this work. B, Results of Boolean dynamics, linear attractors were averaged. C, The occurrence of attractors among all Boolean equation variations. D, Total size of each attractor in all attractors combined. E, An overview of the attractors for all Boolean variations. The color of the circle represents the attractor type (red: linear attractor; blue: point attractor). Aera of the circle represents the attractor size. Each attractor of each Boolean variant shows the same transparency, thus the less transparency represents the higher occurrence. _p: point attractor; _l: linear attractor.

### Bifurcations observed with evolving Hill coefficient

Complex dynamic behavior, like transition states and bifurcation, cannot be obtained in Boolean dynamics(Toulouse et al. 2005; P. Ao et al. 2010; Yuan et al. 2017). Thus, we turn our attention to the ordinary differential equation (ODE) model. In this work, we used a double-normalized Hill equation which has been described in detail previously(Toulouse et al. 2005; Yuan et al. 2017). The details of the direct interactions among genes in the current network are still unclear, let alone their dynamic parameters. In order to extract insights from such limited details, a set of dimensionless mesoscopic parameters, h and k, were used in this work.

We chose h=3, k=10 as the first trial of the parameter set (the vanilla set) based on our previous modelling experience(Yuan et al. 2017). Under this set, we obtained three fixed points, with two stable ones (stable state) and an unstable one (transition state) (Figure 2. A, B). The expression patterns of two stable states (Figure 2. A, B; vanilla_sta1, vanilla_sta2) consist with the point attractors in Boolean Dynamics (Figure 2. A, B; Boole_p1, Boole_p2), and the transition state also consists with the average of all the linear attractors in the Boolean dynamics with the unaltered functions (Figure 2. A, B; Boole_li_allme: the average of Boole_li_7 and Boole_li_18 in Figure 1. B).

**Figure 2.**
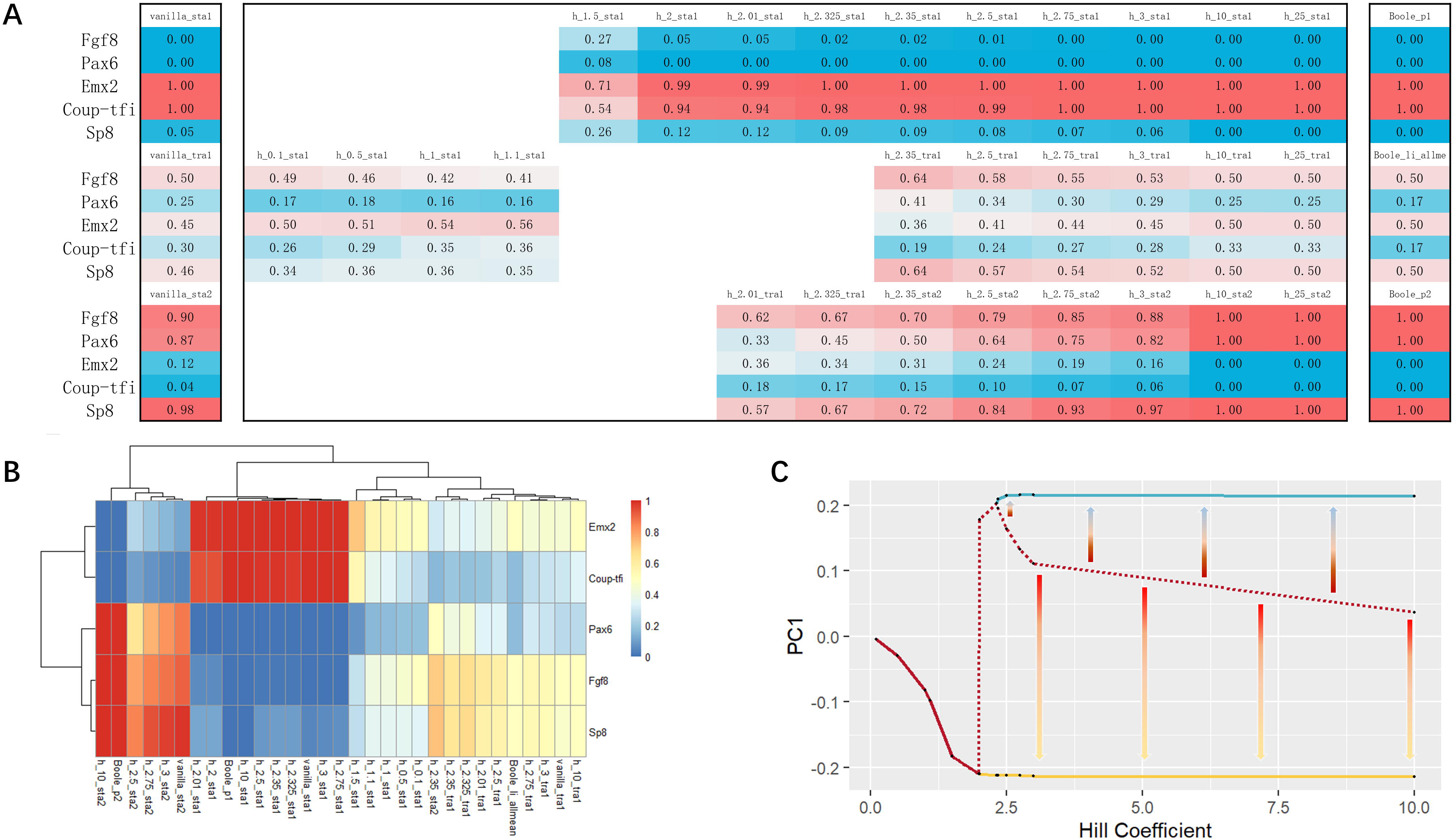
Bifurcations occur when exploring the parameter space. A, A Numeric way of showing the bifurcation. All fixed points under all parameter h we have been tried were displayed in this picture, the first column on the left is under vanilla parameter set (h=3, k=10), the rightest column is the results of the unaltered Boolean functions (h→+∞), and others are arranged from small to large for h. B, Heatmap of all ODE fixed points demonstrated after hierarchical clustering. C, Bifurcations occur with the growth of h. Arrows point to the stable states from the transition state. sta: stable state; tra: transition state

In a nonlinear system, bifurcations may occur as parameters change. Oster and Alberch argued that mathematical bifurcations could be an important mechanism behind development and evolution 40 years ago(Oster and Alberch 1982; François 2012). We then tried to explore the parameter space in order to find bifurcations.

Fitting an elephant with a wiggling trunk is not our purpose, therefore the number of arbitrary parameters needs to be reduced at first. The number of the arbitrary parameters in this model equals 20 (2n, n = edge number). Given that the details of these dynamical parameters are not clear, and genes in the core network level should be considered equally important, as we argued before(Ping Ao et al. 2008; Yuan et al. 2017), thus we took the same parameters for all edges, therefore reducing the arbitrary parameters number from 20 to 2. Considering the concentration/function of genes already normalized into [0, 1], in order to center the S shape part of the sigmoid function, we let k = 2^h, therefore reducing the number of arbitrary parameters to 1.

Increasing h (h=10, 25; k=2^h) results in two stable states and one transition state, which is similar with the dynamic behavior under the vanilla parameter set (Figure 2. A, B). Since the Hill equation is equivalent to its Boolean from h takes the limit, we expected the dynamic behavior will not change significantly with further increased h.

Then we explored the parameter space with smaller h, and discovered two bifurcations (2<h<2.01; 2.325<h<2.35; k=2^h). When h<2, there is only one stable state, which has a pattern similar with the transition state with the vanilla parameter set. The pattern gradually evolves from the vanilla transition state-like pattern to a vanilla stable state 2-like pattern with increasing h. Passing over the first bifurcation (2<h<2.01), a transition state emerges, which then evolves to a vanilla stable state 1-like pattern (2.01<h<2.325). The second bifurcation (2.325<h<2.35) gives birth to the second stable state (Figure 2. C).

### Evolution of the telencephalon anterior-posterior patterning by bifurcations of the network dynamic behavior

Surprisingly, the dynamic behavior of this network is highly consistent with the facts we know about the evolution of telencephalon.

A telencephalon-like structure has been identified recently in amphioxus with a low level of Pax4/6 and a high level of EmxA and EmxB(Benito-Gutiérrez et al. 2021). Only one stable state exists when weak or negative cooperativity among this network (0<h<2.01) with a low level of Pax6 and intermediate-high level of Emx2, which consists with the expression pattern of the telencephalon-like structure identified in amphioxus. The expression pattern of this solitary stable state gradually evolves to a posterior telencephalon-like pattern with the increasing h.

With a further increased h (2.01<h<2.325), the first bifurcation occurs, giving birth to a stable state and a transition state. The expression pattern of the stable state consists with the posterior part of the telencephalon, and the expression pattern of the popping-out transition state is similar with the anterior part of the telencephalon. We don’t know which organism would eventually end in development at this stage. Still, in the development of the vertebrate embryonic central nervous system, the telencephalon first appears near the anterior end of prosencephalon(Briscoe and Ragsdale 2019).

This model results in two stable states consisting with telencephalon anterior-posterior patterning in vertebrates after the second bifurcation (h≥2.35); and one transition state which shares the similarity in expression pattern with the telencephalon-like structure in amphioxus as well as the sable state when weak or negative cooperativity among this network, reminds us of the biogenetic law(Levit et al. 2022).

We did not observe any significant variation in the dynamic behavior of this network after the second bifurcation (h≥2.35). On the biological side, no matter how complex the brain evolves, the binary patterning of pallium/subpallium or olfactory bulb/olfactory cortex can always be observed in the vertebrate telencephalon(Briscoe and Ragsdale 2019).

### Predictions

Based on the results above, we can make the following predictions:

1. The topological structure of the core endogenous network that determines the telencephalon anterior-posterior patterning has already evolved in the ancestral chordate 550 million years ago;
2. The expression pattern of the telencephalon-like structure in such ancestral chordate 550 million years ago and in the Amphioxus today consists with the stable states we obtained before the first bifurcation (0<h≤2);
3. An ancestral organism with primitive telencephalon structure would undergo an expression pattern consistent with the anterior part of the vertebrate telencephalon and the transition state we obtained under the parameter range of 2.01<h<2.325 during development, then would mature in the expression pattern consistent with the posterior part of the vertebrate telencephalon and the stable state we obtained under this parameter range of 2.01<h<2.325.
4. Experiments that reduce the cooperative effect in the network described above in vertebrates, series of events: the vanishing of the anterior-posterior patterning of the telencephalon; a structure with the expression pattern consists with the posterior part of the telencephalon; a structure with the expression pattern consists with the Amphioxus telencephalon-like structure would be observed.

## Conclusion

The current results display such a picture:

The ancestral chordate 550 million years ago evolved a 5-node core endogenous network described in this work with weak or even negative cooperative effect, which determines a telencephalon-like structure with a similar expression pattern with the developing telencephalon during early embryonic stage in the vertebrate, but without the anterior-posterior patterning;

Between 550 million to 500 million years ago, new genes emerged from the genome duplication and mutations. As a result, the cooperative effect in the 5-node core endogenous network increased so that a primitive telencephalon structure gradually evolved to the expression pattern of the posterior part of the vertebrate telencephalon, and during development, it would undergo an expression pattern similar to the anterior part of the vertebrate telencephalon;

With the further increase of the cooperative effect during evolution, a telencephalon with the anterior-posterior patterning emerged, which like the biogenetic law concluded, would undergo a development stage with an expression pattern similar to that of the ancestral telencephalon-like structure.

In a way, we have drawn an answer to how did the telencephalon anterior-posterior patterning evolve quantitively in a first principle manner.

## Discussion

For the evolution of the telencephalon anterior-posterior patterning, we found that such a 5-node network alone is sufficient to draw an answer, but it is not enough for the evolution of the whole central nervous system, let alone the entire vertebrate body plan. The network described in this work can be taken as a subnetwork(Wang et al. 2018), which has the potential to expand into the global decision-making network for the evolution and development of the central nervous system or even the entire vertebrate body plan shaped by evolution.

Preliminary evidence suggest the hypothesis that the network described in this work is conserved during evolution from pre-vertebrate chordate to vertebrate is acceptable, the results obtained by Giacomantonio and Goodhill(Giacomantonio and Goodhill 2010; 2014) and the results described in this work combined further validated this hypothesis at the computational level. Nonetheless, the efforts on the experiments’ side to validate the hypothesis are still indispensable.

As the knowledge of the real dynamics parameters is little, if not none, we used a set of dimensionless mesoscopic parameters and made the parameter sets equal at the core network level based on a set of working hypotheses postulated before(Ping Ao et al. 2008; Yuan et al. 2017). This work showed that useful predictions can still be drawn even with such little knowledge. But exploring the range of these parameters in real biological systems is still a non-negligible task.

In response to the growing emphasis on theory-experiment coupling in life science recently(MacArthur 2023; Nurse 2021; Smaldino 2019; Prokop 2022; Ping Ao 2021), our work showed that the combination of existing experimental facts as well as the aforementioned mathematical and theoretical framework, although not perfect yet, has been able to give birth to new knowledge and verifiable predictions.

## Methods

### Boolean dynamics

For the unaltered function set, we used “AND” to represent the interaction of activation, “NOT” to represent the interaction of inhibition:

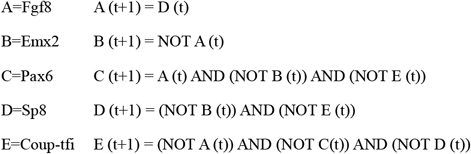

For the Boolean variations, an agent affected by 1, 2, 3 agents will respectively result in 1, 2, 8 equations in this work. Thus, we generated 1*1*8*2*8=128 sets of functions. All variants are presented in the supplemental table.

We iterated the function set with exhausted all initial vectors until stable, then collected and counted attractors, attractor types, and basin of each attractor size. These data are presented in the supplemental table in detail.

### ODE

The ODE functions we used are as follows:

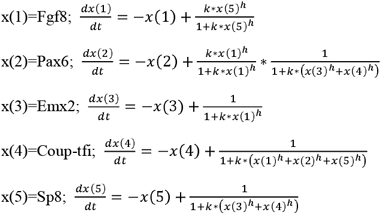

We used Newton’s iteration to obtain fixed points; determined the stability of the fixed point by the real part of the eigenvalue of its Jacobian matrix; used Euler’s method to verify the stable fixed points parallelly.

For the vanilla parameter set (n=3, k=10), we first used 1000 random initial vectors, followed by 100000 random initial vectors to verify the integrity of the results. The results showed that 1000 random initial vectors were sufficient to obtain complete results, so 1000 random initial vectors were used to explore the parameter space.

The connection between transition states and stable states was obtained by perturbation on the transition state with a small random vector; in this work, the number of such small random vectors is 100.

## Contribution

**Design: Ao, Yao;**

**Execution: Sun, Yao, Xiong, Su;**

**Drafting: Yao, Sun, Zhu;**

**Figure: Yao, Sun, Xiong;**

**Revising: Sun, Yao, Xiong, Su, Zhu, Ao.**

## Acknowledgements

This study was supported by grants from the National Natural Science Foundation of China (No. 16Z103060007).

We thank Prof. Wan in the College of Life Sciences, Inner Mongolia Agricultural University for the discussion, Dr. Tang in the Institute of Neurology, Chinese Academy of Sciences and his wife Ms. Cai for the inspiration.

We thank Shanghai Nanobubble Technology Co., LTD. for their assistance in publishing this article.

